# Accurate and efficient discretisations for stochastic models providing near agent-based spatial resolution at low computational cost

**DOI:** 10.1101/686030

**Authors:** Nabil T. Fadai, Ruth E. Baker, Matthew J. Simpson

**Affiliations:** School of Mathematical Sciences, Queensland University of Technology, Brisbane, Queensland 4001, Australia; Mathematical Institute, University of Oxford, Oxford, OX2 6GG, United Kingdom

**Keywords:** compartment based model, lattice based model, proliferation assay, scratch assay, crowding effects, volume exclusion

## Abstract

Understanding how cells proliferate, migrate, and die in various environments is essential in determining how organisms develop and repair themselves. Continuum mathematical models, such as the logistic equation and the Fisher-Kolmogorov equation, can describe the global characteristics observed in commonly-used cell biology assays, such as proliferation and scratch assays. However, these continuum models do not account for single-cell-level mechanics observed in high-throughput experiments. Mathematical modelling frameworks that represent individual cells, often called agent-based models, can successfully describe key single-cell-level features of these assays, but are computationally infeasible when dealing with large populations. In this work, we propose an agent-based model with crowding effects that is computationally efficient and matches the logistic and Fisher-Kolmogorov equations in parameter regimes relevant to proliferation and scratch assays, respectively. This stochastic agent-based model allows multiple agents to be contained within compartments on an underlying lattice, thereby reducing the computational storage compared to existing agent-based models that allow one agent per site only. We propose a systematic method to determine a suitable compartment size. Implementing this compartment-based model with this compartment size provides a balance between computational storage, local resolution of agent behaviour, and agreement with classical continuum descriptions.

## 1 Introduction

Cell-level processes, including proliferation, death, and migration, drive tissue-level processes during regeneration, development and repair [1–3]. Traditionally, mathematical models of development and repair account for such tissue-level processes by modelling the cell population density with ordinary differential equations (ODEs) [4–7] and partial differential equations (PDEs) [8–10]. These continuum descriptions can be parametrised to predict the cell population density growth profile in commonly-used cell biology assays, such as *proliferation assays* [5–7, 11, 12] and *scratch assays* [8–10, 13, 14]. In a proliferation assay (Figure 1), cells are seeded uniformly on a two-dimensional substrate. Due to this initial placement of cells (Figure 1(a),(c)), there are no macroscopic spatial gradients in cell population density at the beginning of the experiment. As the experiment proceeds (Figure 1(b),(d)), individual cells undergo movement and proliferation events, with the net result being a gradual increase in the density of the monolayer towards some maximum carrying capacity density. A common mathematical model to describe these proliferation assays is the logistic equation [4–7],

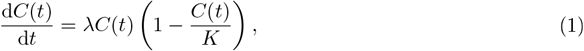

where *C*(*t*) is the cell population density at time *t* ≥ 0, λ > 0 is the cell proliferation rate, and *K* > 0 is the carrying capacity. In a scratch assay (Figure 2), a uniform scratch is made in a cell monolayer and observations are made of the time-dependent movement of the resulting fronts of cells. As the initial scratch creates macroscopic spatial variation in the monolayer, the cell density evolves in both time and space. A common mathematical model to describe these scratch assays is the Fisher-Kolmogorov equation [8, 9, 13–16],

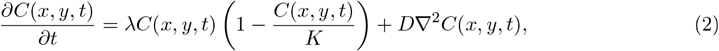

where *C*(*x*, *y*, *t*) is the cell density at time *t* and position (*x*, *y*), and *D* > 0 is the diffusivity of the cell density.

**Figure 1:**
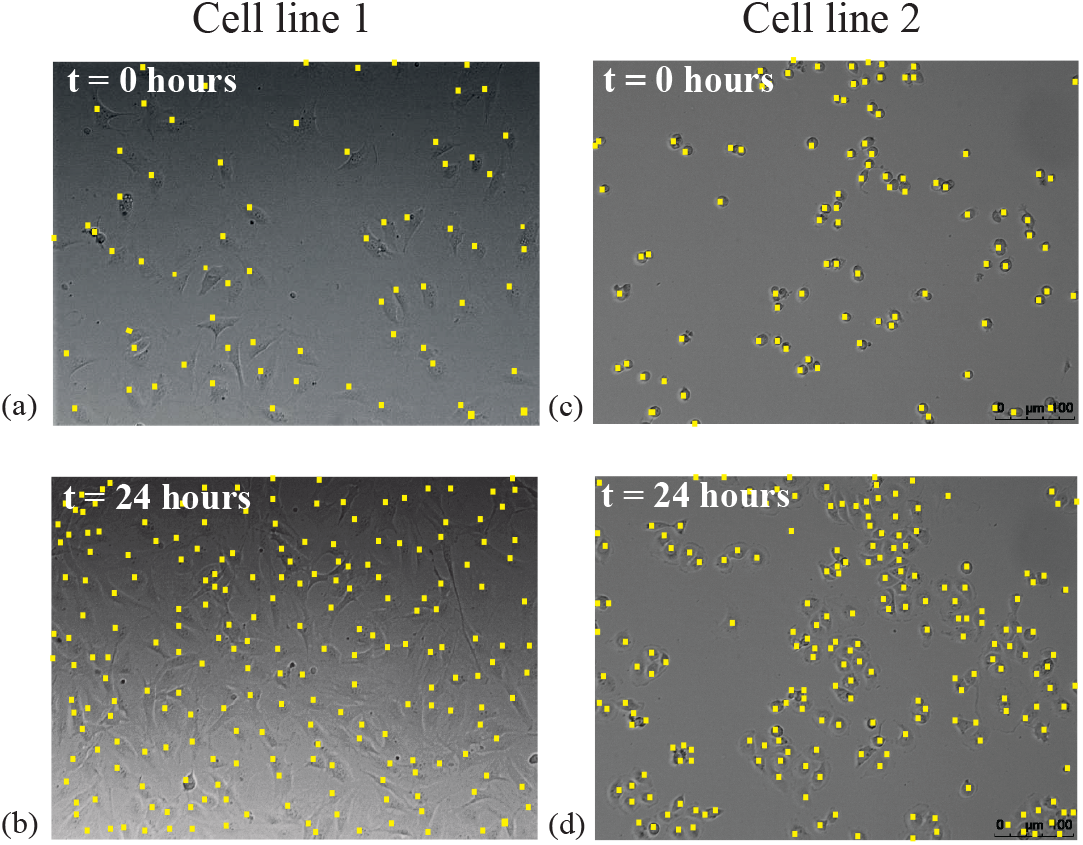
Images of cell proliferation assays. Images in (a)–(b) show a cell proliferation assay with epithelial 3T3 fibroblast cells (cell line 1) [17], while images in (c)–(d) show a cell proliferation assay with mesenchymal MDA-MB-231 breast cancer cells (cell line 2) [18]. The location of cells are highlighted with a yellow marker. The size of images is 640 μm × 480 μm. All images reproduced with permission from [6].

**Figure 2:**
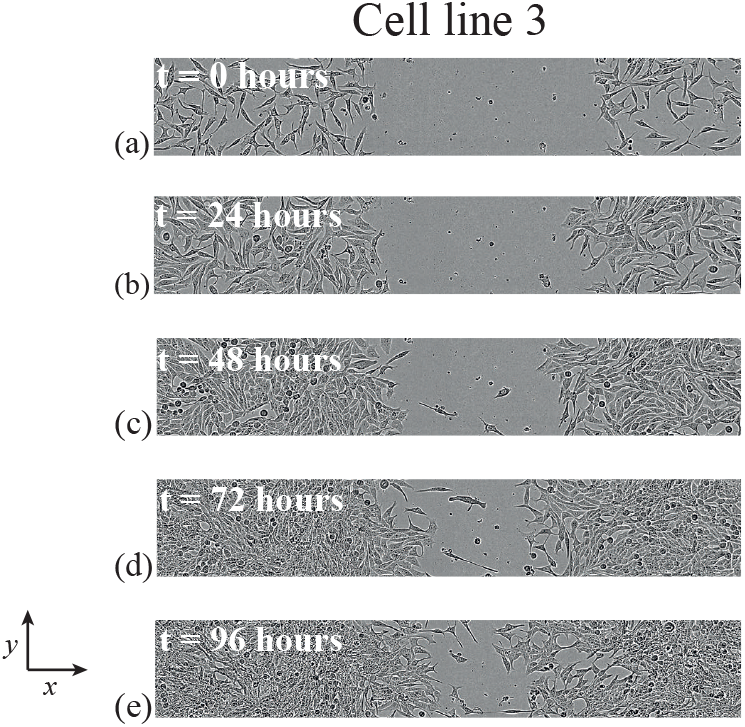
Images of a scratch assay. Images in (a)–(e) show the progression of a scratch assay performed with epithelial C4-2B prostate cancer cells (cell line 3) [19]. The size of images is 1800 μm × 300 μm. The fronts of cells move towards the centre of the initially vacant region as time progresses. All images reproduced with permission from [14].

While Equations (1)–(2) can match the evolution of cell population densities in experiments [8], these models focus exclusively on the characteristics of the *global* cell population. However, recent technology [9, 14, 20, 21] has made it possible to perform these assays in a high-throughput fashion, allowing hundreds of identically-prepared proliferation or scratch assays to be simultaneously performed, as well as to collect single-cell-level data from these assays. With the availability of single-cell-level data, including real-time tracking of cells [20, 21], different types of mathematical models that focus on cell tracking and single-cell-level mechanics are desirable. This modelling approach is especially important since cell-tracking technology is not always accurate, especially in experiments where the cell density is high [22, 23]. A convenient way to model individual cells is in a stochastic mathematical framework; these models are often called *stochastic agent-based models* [5, 7, 16, 24, 25], whereby cells are modelled as *agents*, often constrained to an underlying lattice.

A common spatial discretisation for stochastic agent-based models [5, 7, 16, 24, 25] involves choosing the lattice spacing to be equal to a typical cell diameter. This spatial discretisation is a natural choice when agent-based models include crowding effects, often referred to as an *exclusion process*, since lattice sites are limited to binary occupancy of a single agent. However, employing these models with this level of local resolution can be computationally infeasible for large number of agents [26], whereas the computational storage of the analagous continuum model description is independent of the number of agents. This demand of large computational storage motivates us to consider other spatial discretisations. For instance, one could instead choose the lattice spacing to be *m* times the size of a typical cell diameter, where *m* > 1 is an integer, allowing multiple agents to be accommodated within each lattice *compartment*. An immediate consequence of choosing *m* > 1 is a reduction in computational storage, since the computational memory requirement reduces from 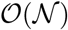, where 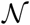 is the number of agents, to 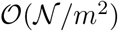, where 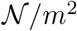 is the number of compartments. However, this reduction in computational storage comes at the cost of losing local agent resolution, so the key question is: how do we choose this lattice compartment size to capture local agent dynamics while still being computationally efficient?

In this work, we propose a modification to previous descriptions of stochastic agent-based models on lattices (e.g. [5, 7, 16]) to: (i) allow for computationally efficient simulations, (ii) capture local agent dynamics, and (iii) provide reliable agreement of the average agent density to traditional continuum model descriptions. This *Compartment-Based Model* (CBM), discretises space using lattice compartments with *m* > 1 (Figure 3). The CBM also encodes additional biologically inspired features, such as crowding effects, whereby potential movement and proliferation events cannot occur if the target compartment is fully occupied with agents. Inclusion of crowding distinguishes the CBM from other agent-based models that describe reaction-diffusion processes [27–34] and provides additional biological realism, since many experimental observations confirm that crowding effects are very important in proliferation assays and scratch assays [1, 3, 5, 6, 11].

**Figure 3:**
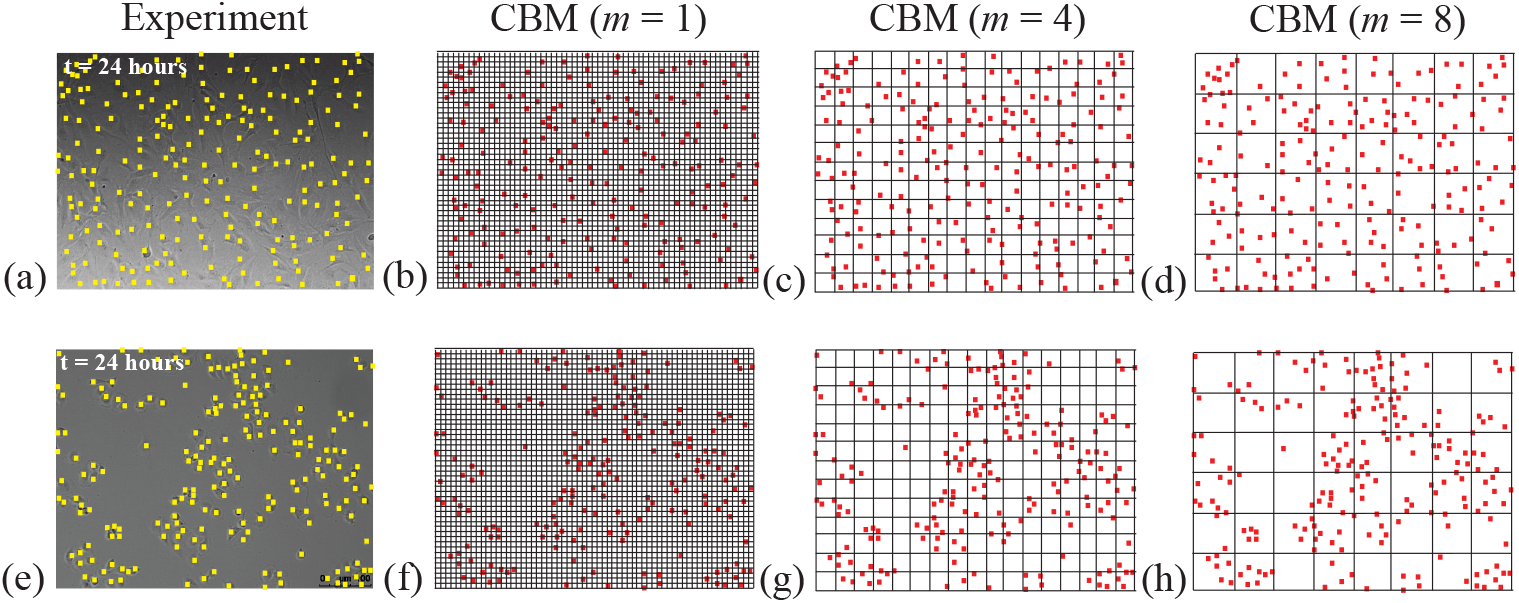
Choices of compartment size in the CBM. Images of proliferation assays using cell lines 1 and 2, from Figure 1(c,d), are shown in (a,e), respectively. Images in (b)–(d) show various discretisations of the arrangement of cells in (a) with *m* = 1, 4, 8, respectively. Images in (f)–(h) show various discretisations of the arrangement of cells in (e) with *m* = 1, 4, 8, respectively.

We compare the CBM with ODE and PDE descriptions, since traditional continuum models provide well-understood explicit solutions to key features of experiments, such as the temporal evolution of the agent density [5–8, 11, 12, 16, 24, 25]. We do this by examining the *continuum limit* of the CBM, which describes the salient features of the CBM in the limit when the number of lattice sites is large [5, 7, 11, 16, 24, 25]. We explore how the predicted average density of agents changes for different compartment sizes. In particular, with larger *m*, the CBM allows us to retain the use of traditional continuum descriptions when cell clustering is either absent (Figure 1(a),(b)) or present (Figure 1(c),(d)). Previous examination of stochastic agent-based models [5, 7, 16] reveals that when the proliferation-to-motility rate ratio is not sufficiently small, clustering develops and the resulting agent density profile no longer matches the solution of the continuum limit when *m* =1 (Equations (1)–(2)). However, the CBM avoids this disagreement by using a sufficiently large compartment size. We show that, for a suitable choice of *m*, the average agent density determined by the CBM agrees well with the solution of the continuum limit and provides a balance between computational storage and local agent information.

## 2 Model

We begin by presenting the CBM on a two-dimensional square lattice (Figure 3) to describe simulations of both proliferation and scratch assays. Additional results (Supplementary Material) demonstrate how the CBM generalises to three-dimensional lattices that are relevant for three-dimensional assays.

### 2.1 Lattice discretisation

As we focus on employing the CBM to describe two-dimensional assays (Figure 3), we consider ways to discretise an 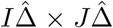 rectangle with a square lattice. Here, 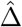 is a typical cell diameter (20-25 μm, [6]), implying that there can be most 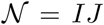 agents on the lattice under square packing. To compare different cell lines with different cell diameters, we non-dimensionalise the lattice to have unit length spacing by setting 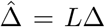, where Δ = 1 and *L* is the cell diameter [5, 6, 25]. We focus on non-dimensional lattices and represent the location of the top right corner of each site in Cartesian co-ordinates as (*x_i_*, *y_j_*) = (*i*Δ, *j*Δ), where *i* = 1, …, *I* and *j* = 1, …, *J*.

The CBM discretises this underlying lattice into *compartments*. These compartments, of size *m*Δ × *m*Δ, can contain up to *m*^2^ agents. To ensure that the dimensions of the original 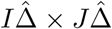 rectangle remain consistent under this discretisation, we have *X* = *I/m* compartments in the *x*-direction and *Y* = *J/m* compartments in the *y*-direction. We index each *m* × *m* compartment with co-ordinates (*x_i_*, *y_j_*) = (*im*Δ, *jm*Δ), where *i* = 1, …, *X* and *j* = 1, …, *Y*. While the total number of compartments in the CBM is 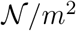, the maximum number of agents on the lattice stays at 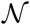.

The CBM is an extension of previous agent-based models with crowding [5, 7, 16], in which each compartment can be occupied by at most one agent (*m* = 1). Here, the distinguishing feature of the CBM is the fact that each compartment can be occupied by more than one agent when *m* > 1 and the key question is how we choose *m* to reduce computational storage while retaining sufficient local resolution. In Figure 3, snapshots of proliferation assays are used to motivate the choice of *m*. As previously mentioned, the CBM with *m* = 1 (Figure 3(b,f)) demands significant computational memory for large numbers of agents. Contrastingly, employing the CBM with *m* > 1 (Figure 3(c,d,g,h)) reduces the computational storage. Both of these advantages, along with a systematic method of determining an appropriate compartment size, will be discussed in Section 3.

### 2.2 The Compartment-Based Model (CBM)

Since crowding effects are important in cell biology assays [5, 7, 9, 16, 25], the CBM is an exclusion process, as both movement and proliferation processes may only take place if the target compartment (i.e., either the same or an adjacent compartment of the agent undergoing these processes), has sufficient space to accommodate potential motility and movement events. For lattice-based models, such as the CBM, the excluded volume is the volume occupied by agents; however, this equivalence is not true for lattice-free agent-based models [35, 36]. At any time, a randomly chosen isolated agent has a transition rate *r_m_* per unit time of moving (either within the same compartment or to an adjacent compartment), a proliferation rate *r_p_* per unit time of giving rise to another agent (placed either in the same or an adjacent compartment), and a death rate *r_d_* per unit time (agent is removed). We assume that an agent is equally likely to be found at any particular location within a particular compartment; this is also known as a *well-mixed* assumption. The probability of an isolated agent attempting to move or proliferate to an adjacent compartment, rather than remain within the same compartment, is 1/*m*. This probability can be interpreted as the number of configurations, for an agent placed inside a well-mixed *m* × *m* compartment, that result in the agent moving or proliferating into an adjacent compartment, 4*m*, divided by the total number of configurations, 4*m*^2^. The probability that there is sufficient space available in the compartment selected for the agent to move or proliferate into is 1 – *N*/*m*^2^, where *N* is the number of agents in this compartment. We implement reflecting conditions along all boundaries of the lattice, which models zero net flux of cells into or out of the domain [9, 21, 37]. The initial placement of agents is discussed further in Sections 3.1 and 3.2. Using the Gillespie algorithm [38], we simulate the evolution of agents as a function of time and space using Algorithm 1.

To quantify data from the CBM, we introduce appropriate notation. To describe a proliferation assay, we define *Q_m_*(*t*) as the total number of agents on the lattice discretised with a compartment size *m* at time *t* and from a single realisation of the CBM. When comparing data from the CBM with the continuum limit description for a proliferation assay, we average data from the CBM using

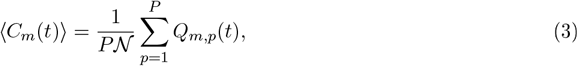

where *Q_m,p_*(*t*) is the *p*th identically prepared realisation of *Q_m_*(*t*) and *P* is the total number of identically-prepared realisations. To describe a scratch assay, we define *Q_m_*(*x_i_*,*y_j_*, *t*) as the number of agents in each *compartment* of size *m* located at co-ordinates (*x_i_*,*y_j_*), at time *t*, from a single realisation of the CBM. Motivated by the scratch assays in Figure 2, we examine spatially dependent initial conditions that are approximately uniform in the *y*-direction [9, 21, 37, 39]. To compare data from the CBM with the continuum limit description for a scratch assay with *y*-independent initial conditions, we average data in the *y*-direction alone using

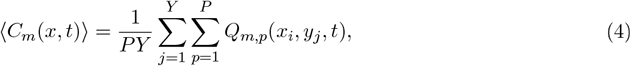

where *Q_m,p_*(*x_i_*,*y_j_*, *t*) denotes the number of agents in a compartment at position (*x_i_*,*y_j_*) in the *p*th identically prepared realisation.

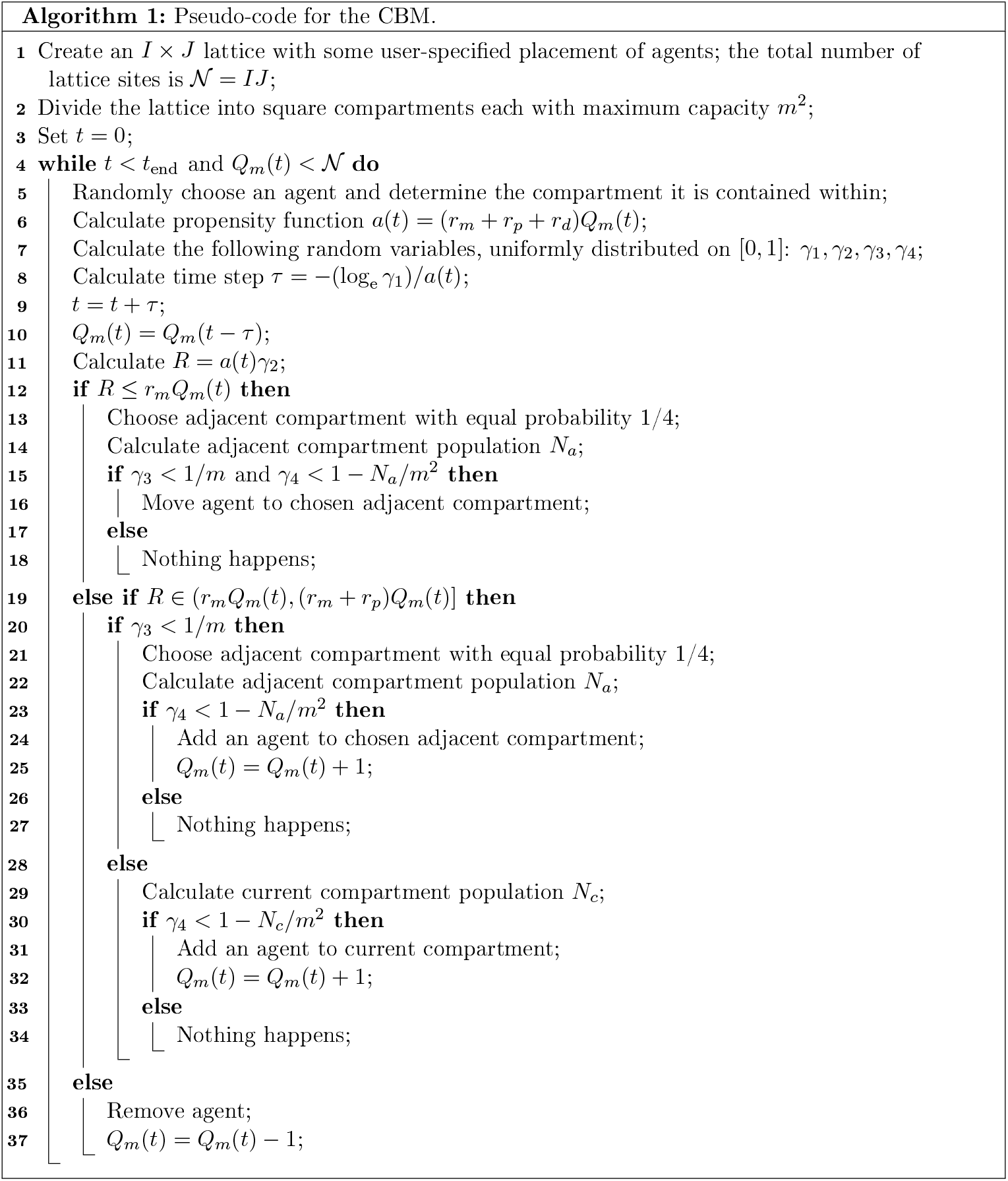

### 2.3 Continuum limit

An aim of this work is to formulate and implement the CBM so that the averaged data from this stochastic model is consistent with commonly-used traditional continuum descriptions. Following [24, 25, 28], we examine the limit when the number of sites is large. In this limit, we can arrive at mathematical descriptions of the time-dependent average density by constructing approximate conservation statements and taking appropriate limits [24, 25, 28]. Furthermore, by assuming that the occupancy status of lattice sites are independent (normally referred to as the *mean-field approximation* [5, 7, 11, 16, 24, 25]), the continuum limit of the CBM with *m* = 1 [24, 25] is the two-dimensional analogue of the Fisher-Kolmogorov equation (Equation (2)) on the domain [1, *I*] × [1, *J*], where λ = *r_p_* – *r_d_*, *K* = 1 – *r_d_*/*r_p_*, and *D* = *r_m_*Δ^2^/4. For the CBM, we require that Equation (2) be modified for *m* > 1; further details of these modifications appear in Section 3.

In previous examination of continuum limits of stochastic agent-based models with crowding, such as the CBM with *m* = 1 [5, 25], agreement between the solution of Equations (1)–(2) and the averaged agent density from the stochastic model requires *r_p_*/*r_m_* ≪ 1 (see Figure 4(a)–(d)). This agreement occurs because the mean-field approximation is valid in this parameter regime [5, 7, 16]. In Section 3, we will show that, with a suitable choice of *m*, agreement between the solution of the continuum limit and the CBM average agent density is excellent, even when *r_p_*/*r_m_* is not sufficiently small and limit and the CBM average agent density is excellent, even when *r_p_*/*r_m_* is not sufficiently small and clustering is present (Figure 4(e)–(h)).

**Figure 4:**
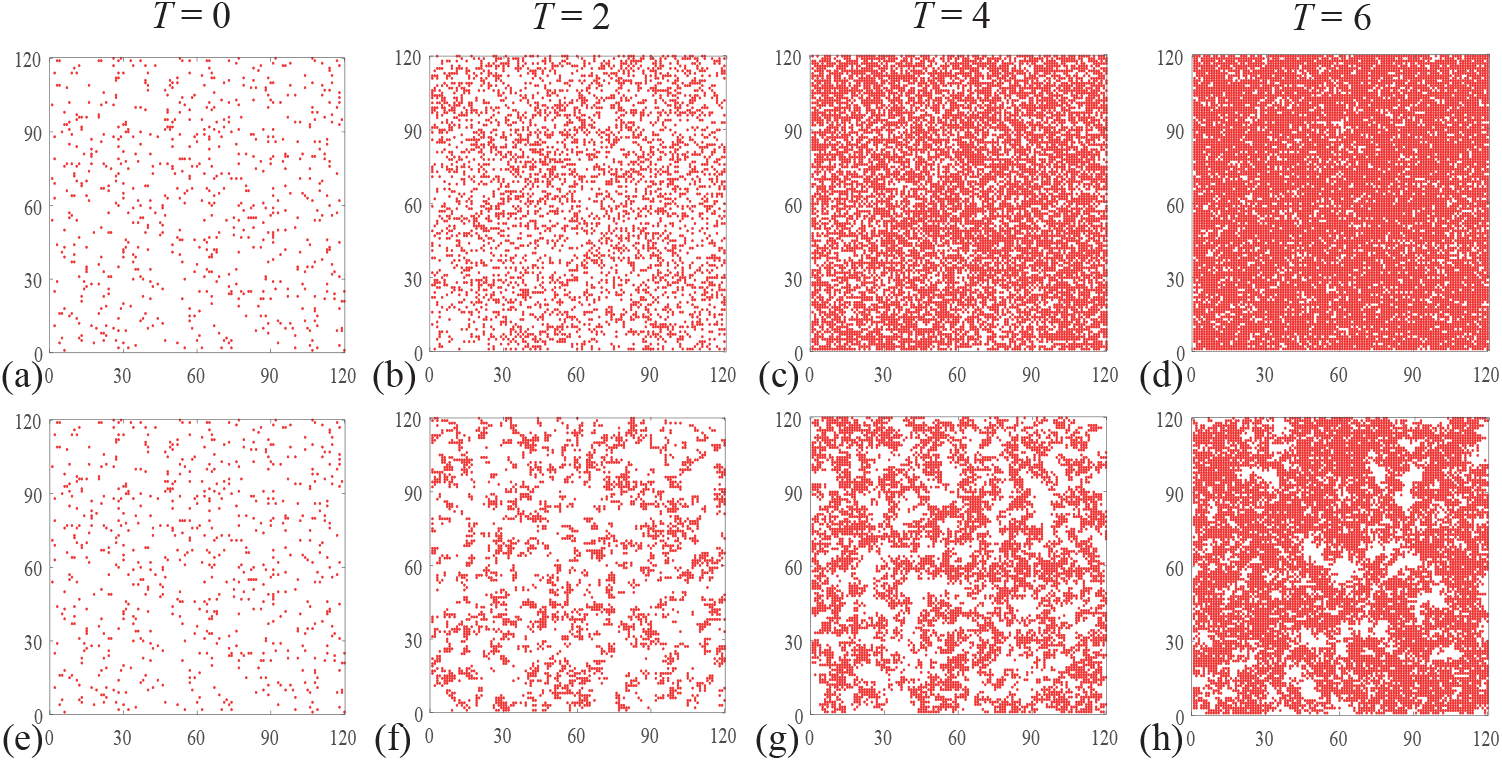
Simulations of two cell proliferation assays using the CBM with *m* = 1. The first simulation, shown in (a)–(d) with *r_m_* = 1 and *r_p_* = 0.01, results in the absence of clusters. The second simulation, shown in (e)–(h) with *r_m_* = 1 and *r_p_* = 1, results in clear cluster formation. In both simulations, each site of the corresponding 120 × 120 lattice is initially populated uniformly at random with probability 0.05. To compare simulations with different proliferation rates, we show results on the non-dimensional timescale *T* = λ*t* = (*r_p_* – *r_d_*)*t*, where *r_d_* = 0.1*r_p_*.

## 3 Results and discussion

It now remains to show how one should choose *m*. Since the CBM only keeps track of the occupancy of compartments, rather than individual agents’ locations, the computational storage decreases as *m* increases. However, this comes at the cost of losing local agent resolution. Consequently, it is important to determine the *minimum* compartment size for which the average agent density of the CBM reasonably matches the solution of the continuum limit.

### 3.1 Simulating cell proliferation assays using the CBM

We begin by employing the CBM in a setting that is appropriate for modelling a cell proliferation assay. Here, we focus on *I* × *I* lattices (i.e., *I* = *J*), so that 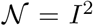. The proliferation assay begins with uniformly seeded agents, with no macroscopic gradients in agent density (Figure 4(a),(e)). In the context of the CBM, each site is initially populated uniformly at random, with some user-specified probability. Because of these translationally invariant initial conditions, the net flux of agents entering and leaving each compartment due to migration is zero and ∇^2^*C* = 0 [9, 21, 37]. Consequently, *C*(*x*, *y*, *t*) simplifies to a function of time only, *C*(*t*), and Equation (2) simplifies to Equation (1), which can be written as

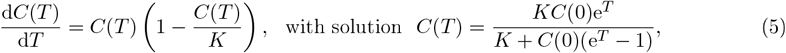

where *T* = λ*t* and *C*(0) is the initial agent density. Previous examination of stochastic models with *m* = 1 and *r_p_*/*r_m_* ≪ 1 [5, 7, 13, 16] reveal that the averaged model data agrees well with the solution of the continuum limit, Equation (5). Under these conditions, pairwise correlations between the occupancy status of lattice sites are negligible. From [5, 7, 13, 16], we note that pairwise correlations are a *local* effect and need only be considered for small distances between agents.

To quantify how far sites must be separated before the correlation in occupancy is negligible, we examine the *correlation function*, *F*(*s*,*T*) [5], using the CBM with *m* = 1. The correlation function is

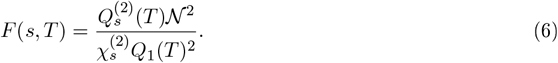

Here, 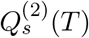 is the number of pairs of agents separated on the underlying *m* = 1 lattice by the Euclidean distance *s* at time 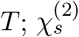 is the number of distinct lattice site pairs separated by a distance *s*. By denoting the discrete two-dimensional Euclidean distance *s* as 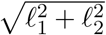, where *ℓ*_1_ = 1, …, *I* and *ℓ*_2_ = 0, …, *ℓ*_1_ to prevent double-counting, we express 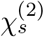 as

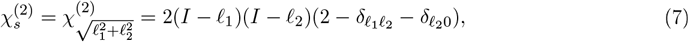

where *δ_ij_* is the Kronecker delta. To simplify notation, we will refer to the set of discrete Euclidean distances separated by 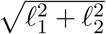 as {*s_k_*}, *k* ≥ 1.

If all sites are uncorrelated, *F* = 1, which is implicitly assumed in deriving the continuum limit descriptions in Section 2.3 [5, 7, 16]. To make an appropriate choice of *m*, we wish to determine the *threshold correlation radius*, 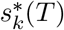, beyond which pairwise correlations are sufficiently negligible:

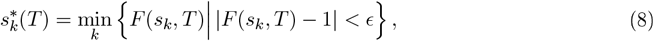

for some user-specified tolerance *ϵ* > 0. Furthermore, we define the *maximal correlation radius* as

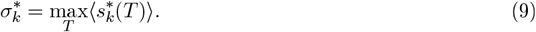

Here, 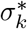 is implicitly a function of *r_m_*, *r_p_*, *r_d_* and *C*(0), and 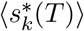 is the average of 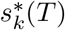 over many realisations of the CBM with *m* = 1.

The maximal correlation radius, 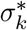, is straightforward to compute, with the advantage that it only relies only on quantities that are available during a typical simulation of the CBM when *m* = 1, for any *r_m_*, *r_p_*, *r_d_*, and *C*(0). We now show how to determine a suitable compartment size *m* from 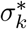. There are many ways to choose the *minimum* compartment size, denoted as *m*\*, from 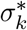, provided that *m*\* is a monotone non-decreasing function in 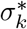 and that *m*\* = 1 when the mean-field approximation is satisfied up to the tolerance *ϵ*. We will consider the function

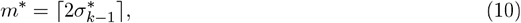

since all significant pairwise correlations are contained within a compartment *diameter* of at least 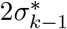. Therefore, any choice of *m* ≥ *m*\* ensures that the mean-field approximation is satisfied up to the tolerance *ϵ*. We note that if the mean-field approximation is satisfied up to the tolerance *ϵ* for all time, then 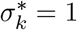, i.e. *k* = 1. From Equation (10) and defining 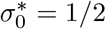, we have that *m*\* = 1 is the minimal compartment size when the mean-field approximation always holds up to tolerance *ϵ*.

By definition, the choice of *ϵ* will influence how large the maximal correlation radius 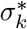 becomes. However, there is good agreement between the average agent density of the CBM with *m* = 1 and the solution of the continuum limit when *r_p_*/*r_m_* ≪ 1 [5, 7, 16], we should choose *ϵ* in such a fashion that 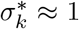 in this parameter regime. Previous results in [5, 7], as well as the Supplementary Material of this work, show that when *F* < 1.5, there is excellent agreement between the CBM (*m* = 1) and the solution of the continuum limit, implying that *ϵ* < 0.5 is sufficiently small. Indeed, as shown in Figure 5(a), employing the CBM in this parameter regime with larger compartment sizes (*m* = 4, 6) does not significantly affect the agreement between 〈*C_m_*(*T*)〉 and *C*(*T*).

**Figure 5:**
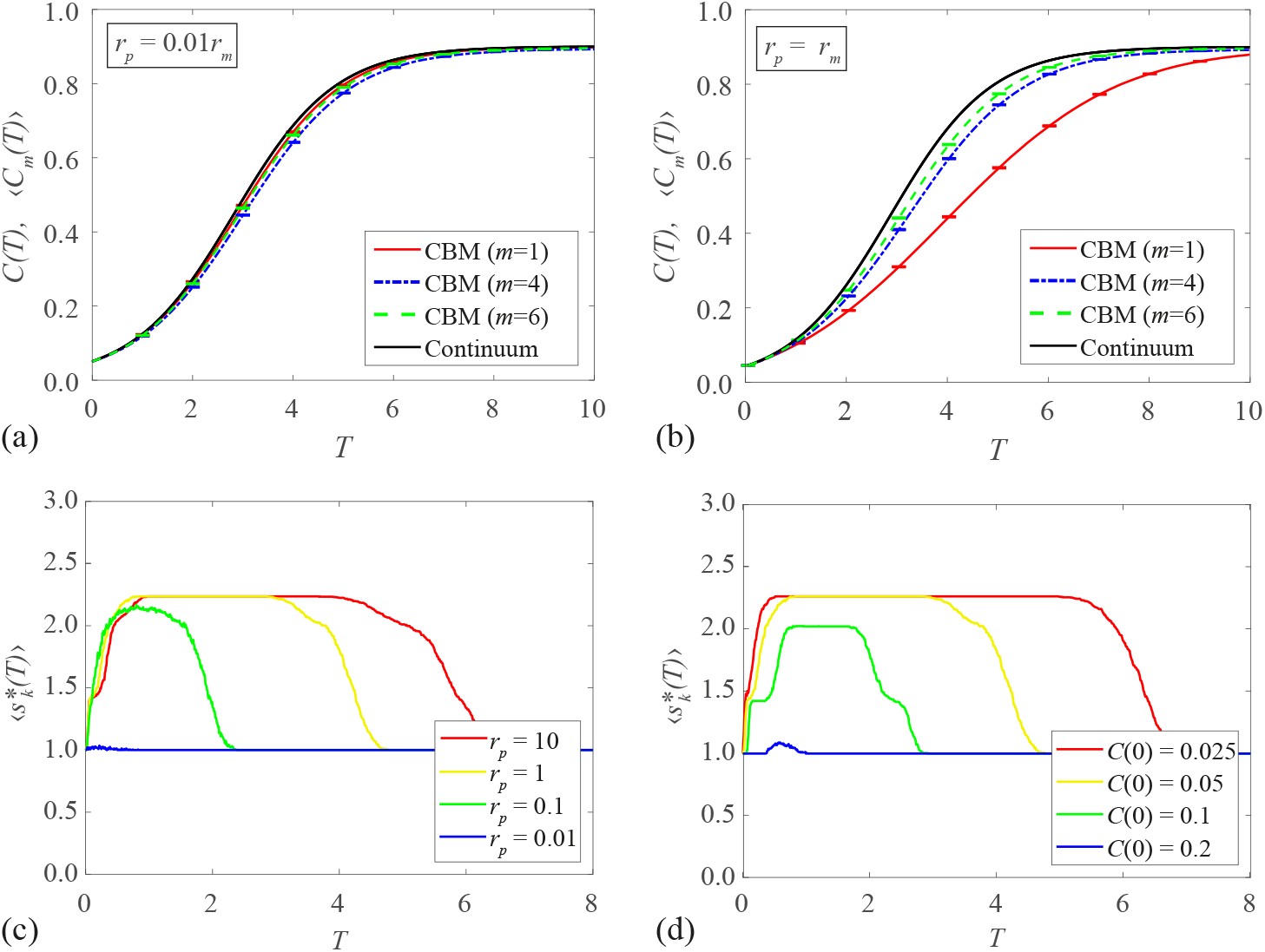
Simulations of a proliferation assay with the CBM. The CBM uses a 120 × 120 lattice, *r_m_* = 1, and *r_d_* = 0.1*r_p_*, while 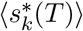, defined in Equation (8), uses the tolerance *ϵ* = 0.3. In all simulations, we use the same initial conditions as in Figure 4. Averaged density data in (a)–(b) is constructed using 100 identically prepared realisations of the CBM. Results are shown for the CBM with *m* = 1, *m* = 4, and *m* = 6, and the continuum limit is given by Equation (5). In (a), *r_p_* = 0.01; in (b), *r_p_* = 1. In both (a) and (b), the maximum standard error (Supplementary Information) is less than 0.002. (c) Comparisons of 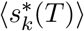 for various choices of *r_p_*. (d) Comparisons of 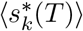 for various choices of *C*(0) with *r_p_* = 1.

Knowing that 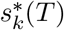 is constructed to provide agreement between the average agent density of the CBM and the solution of the continuum limit where clustering is absent, we now examine parameter regimes where agent clustering is present. As is evident in Figure 4(e)–(h), clusters of agents are visually distinct when *r_p_*/*r_m_* = 1, providing a suitable parameter regime to test how robust the CBM is. Results in Figure 5(b) confirm that the agreement between 〈*C_m_*(*T*)〉 and *C*(*T*) improves as the compartment size *m* increases. Additionally, we note that the threshold correlation distance 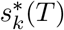 predicts that 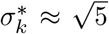 (yellow curves, Figures 5(c)–(d)), implying, from Equation (10), that *m*\* ≈ ⌈2 · 2⌉ = 4 is the minimum compartment size to sufficiently contain pairwise correlations in this parameter regime. Therefore, we do not expect that the average agent density from the CBM would agree with the solution of the continuum limit in this parameter regime for *m* = 1. Nevertheless, a compartment size *m* larger than *m*\* (say, *m* = 6, green curve in Figure 5(b)) is sufficient in obtaining reasonable agreement.

Finally, we examine how 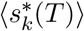 varies with *r_p_*/*r_m_* and *C*(0). Without loss of generality, we set *r_m_* = 1 and examine the influence of *r_p_* and *C*(0) on 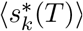. As shown in Figure 5(c), the maximum value of 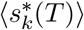, which is 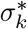, decreases as *r_p_* decreases. This is to be expected; a small *r_p_*/*r_m_* corresponds to parameter regimes where the average agent density of the CBM with *m* = 1 matches the solution of the continuum limit. Furthermore, this same phenomenon happens when *C*(0) is *increased* (Figure 5(d)). Therefore, in parameter regimes where *C*(0) is small, or when *r_p_*/*r_m_* is sufficiently large, *m*\* > 1 and thus disagreement is expected. Therefore, choosing a compartment size *m* > *m*\* in the CBM reduces computational storage requirements, provides better agreement with the solution of the continuum limit when agent clustering is present, and retains local agent behaviour.

### 3.2 Simulating scratch assays using the CBM

Now having demonstrated that, for a suitable *m* and *ϵ*, the average agent density of the CBM agrees with the solution of the continuum limit for cell proliferation assays, we examine how the CBM can be used to describe scratch assays by employing spatially varying initial conditions (Figure 6). We consider simulations of scratch assays where clustering is absent (see cell line 1 in Figure 1, Figure 6(a)–(e)) and simulations of scratch assays where clustering is present (cell line 2 in Figure 1, Figure 6(f)–(j)). To apply the CBM, we must first consider how the diffusion term in Equation (2), *D*∇^2^*C*, changes when varying *m*. Previous examination of the continuum limit of diffusion-only compartment-based models [28, 29] reveals that the jump rates between adjacent compartments scales with 1/*m*^2^, for *m* > 1. However, the models proposed in [28, 29] assume that an isolated agent will always leave its compartment, rather than having non-zero probability to remain within its compartment. Since agents in the CBM attempt to move out of a particular compartment with probability 1/*m*, we divide the scaled diffusivity proposed in [28, 29], *m*^2^*D*, by *m*. Therefore, the diffusivity *D* of the CBM continuum limit with *m* = 1 becomes *mD* for the CBM continuum limit with *m* > 1, and the continuum limit description of CBM simulations of scratch assays can be written as

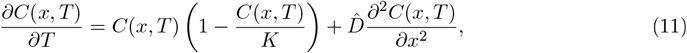

where 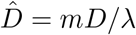. This continuum description is valid when the initial conditions are independent of *y* [39], such as in Figures 2, 6, and 7.

**Figure 6:**
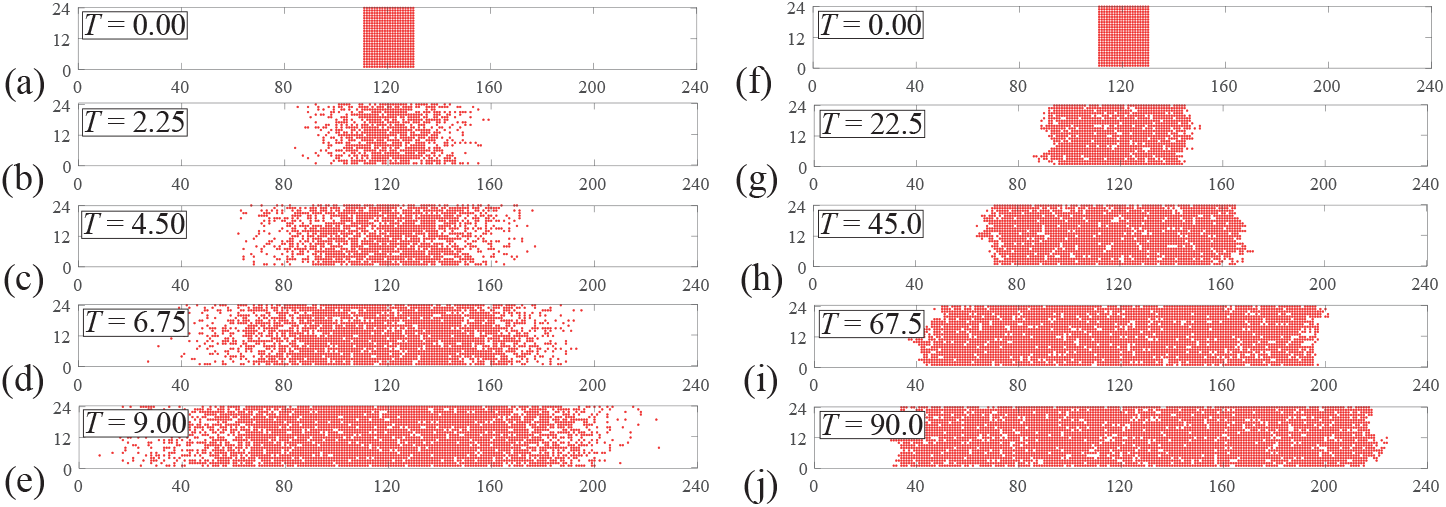
Simulations of two scratch assays using the CBM with *m* = 1. The first simulation, shown in (a)–(e) with *r_m_* = 1 and *r_p_* = 0.01, results in diffuse fronts and the absence of clusters. The second simulation, shown in (f)–(j) with *r_m_* = 1 and *r_p_* = 1, results in clear cluster formation. Both simulations use a 24 × 240 lattice that is initially fully occupied with agents in the region 110 < *x* ≤ 130. In all simulations, *r_d_* = 0.1*r_p_*.

**Figure 7:**
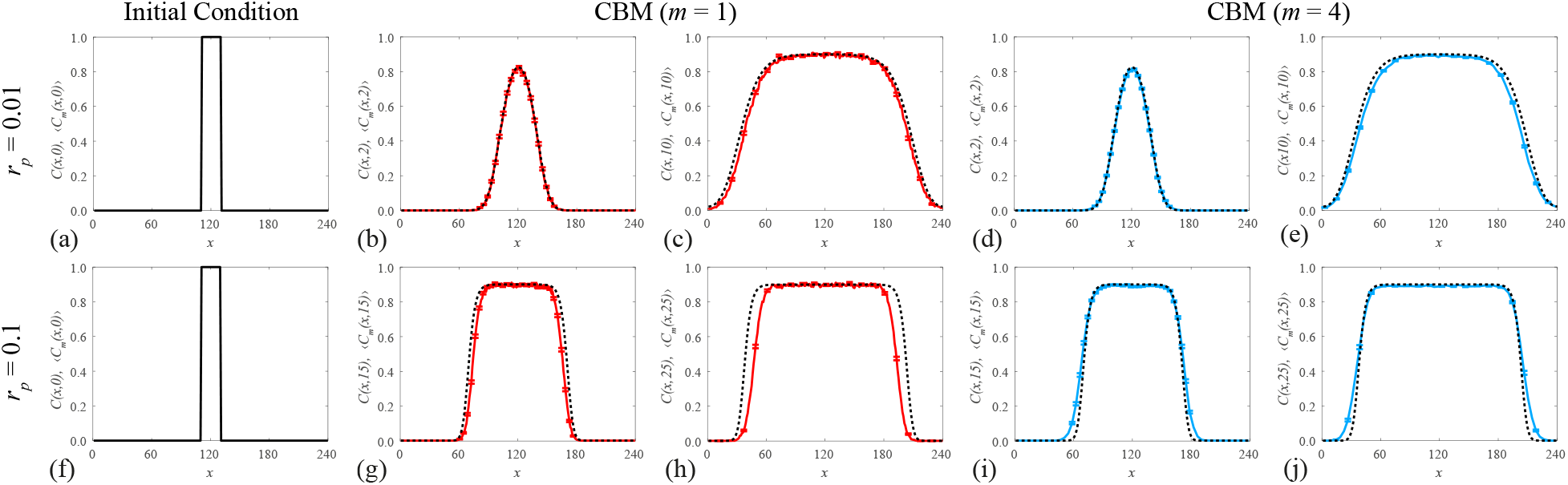
Simulations of a scratch assay with the CBM. Averaged density data from the CBM (solid colour curves) from 100 identically prepared simulations is compared with the solution of the continuum limit, Equation (11) (dashed black curves), for two different scratch assay scenarios. Results are shown for the CBM with compartment size (b,c,g,h) *m* = 1 and (d,e,i,j) *m* = 4. In all simulations, we use a 24 × 240 lattice, with initial conditions shown in (a,f), *r_d_* = 0.1*r_p_*, and *r_m_* = 1/*m* to keep 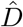 invariant for varying *m*. Top row: *r_p_* = 0.01, with data given at (b,d) *T* = 2 (c,e) *T* = 10. Bottom row: *r_p_* = 0.1, with data given at (g,i) *T* = 15 (h,j) *T* = 25. The maximum standard error (Supplementary Information) is less than 0.012.

Unlike in Section 3.1, it is less obvious how to determine *m*\* when describing scratch assays. This is because the threshold correlation radius (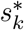 from Section 3.1) will depend on both *T* and *x*, due to the spatially dependent initial conditions. While there are many ways one could determine *m*\* from 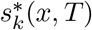, for simplicity, we choose the same *m*\* determined from Equation (10) in Section 3.1. When *r_p_*/*r_m_* ≪ 1 (e.g. *r_p_* = 0.01, Figure 7(a)–(e)), *m*\* = 1 and the average agent density of the CBM, 〈*C_m_*(*x*, *T*)〉, agrees well with the solution of Equation (11) for different compartment sizes and different times. However, for larger proliferation rates (e.g. *r_p_* = 0:1, Figure 7(f)–(j)), the fronts in the CBM become more disperse as *m* increases. While the CBM with *m* = 1 predicts slower moving fronts than the solution of the continuum limit (Figure 7(g,h)), the CBM with an intermediate compartment size (*m* = *m*\* = 4) agrees well with the solution of the continuum limit on the timescale shown in Figure 7(i,j). For larger compartment sizes, the fronts in the CBM are overly disperse and the agreement with the solution of the continuum limit diminishes. These benefits continue to hold for CBM simulations of scratch assays with different initial cell densities (Supplementary Information). Nevertheless, the average agent density of the CBM, 〈*C_m_*(*x*, *T*)〉, can produce qualitatively similar results as the solution of the associated continuum limit for a suitable compartment size, while requiring less computational storage than previously described stochastic agent-based models.

## 4 Conclusions

In this work, we propose a computationally efficient and accurate agent-based model that can be used to simulate two-dimensional cell biology assays. This CBM stems from previous examination of cell proliferation assays, where an initially uniform distribution of biological cells move and proliferate to give rise to a monolayer of cells whose density increases with time. We also apply the CBM to scratch assays, which are prepared by scratching a monolayer of cells and observing the movement of the resulting fronts of cells. The CBM faithfully describes the behaviour of individual cells in these cell biology assays while requiring less computational overhead than other lattice models [5, 7, 16] when modelling large numbers of cells. Furthermore, in parameter regimes that are prone to cell clustering, the average cell density determined by these previous models does not always agree with their continuum description.

We show that the CBM is more computationally efficient than previously proposed exclusion process models on lattices through the discretisation of the underlying lattice into compartments containing multiple agents. These mesoscale compartments in the CBM provide a balance between traditional continuum models, and other agent-based on-lattice models that demand significant computational storage for large numbers of cells. Furthermore, when compartments larger than a threshold size are employed, the CBM agrees well with the continuum description for all physically relevant parameter regimes, including when cell clustering is present. We find that this threshold compartment size is the lattice distance beyond which pairwise correlations of agents are negligible and can be computed directly from the lattice-based model, rather than relying on continuum approximations. Furthermore, this threshold distance is a function of the cell proliferation-to-motility ratio, as well as the initial cell density. We see good agreement between the average agent density of the CBM and the continuum description both in translationally invariant environments (cell proliferation assays) and with spatially dependent initial conditions (scratch assays).

Further extensions to the description of the CBM can be made when comparing to other cell biology experiments. For example, three-dimensional gel proliferation assays describe the proliferation of agents in a three-dimensional material. By considering a three-dimensional setting (see Supplementary Material), we can describe these gel proliferation assays using the CBM. Other kinds of assays observe the chemotactic movement of cells. By biasing the agent movement between compartments (see Supplementary Material), we can describe these chemotactic assays using the CBM. However, the description of the CBM can be further extended to represent additional phenomena present in other biological experiments, including (but not limited to) lattice-free models with crowding, modelling multiple cell types with different proliferation and motility rates, modelling multiple cell types of different sizes, and describing Allee-type dynamics within a single cell type.

## Authors’ Contributions

N.T.F. created the algorithm code, produced Figures 3–7, and carried out the analysis of the results; M.J.S. conceived of the study, designed the study, supervised the study, and helped draft Figures 3–7; R.E.B. participated in the design of the study. N.T.F. wrote the paper, on which all other authors commented and made revisions. All authors gave final approval for publication.

## Acknowledgements

We thank the two anonymous referees for their helpful comments.

