## Supplementary Material for "Accurate and efficient discretisations for stochastic models providing near agent-based spatial resolution at low computational cost"

September 24, 2019

### Contents

**S1 Determining the threshold of the correlation function** **2**

**S2 Three-dimensional simulations** **4**

**S3 Computing the standard error of the mean** **5**

**S4 Varying the initial condition of the scratch assay** **6**

**S5 Biased movement** **8**

**S6 Numerical solution of continuum limit** **10**

### S1 Determining the threshold of the correlation function

In Section 3.1, we note that the choice of $\epsilon$ in the threshold compartment distance, $s_k^*(T)$, determines how close the correlation function, $F(s,T)$, needs to be to unity before pairwise correlations of agents are considered negligible. In other words, the choice of $\epsilon$ determines how far apart agent pairs have to be before they are considered uncorrelated. To determine a suitable choice of $\epsilon$, we focus on two parameter regimes: one where good agreement is seen between the average agent density, $\langle C_1(T)\rangle$, and its continuum limit, $C(T)$, and one where there is poor agreement. In doing so, we can provide a suitable range for $\epsilon$ to use in the definition of $s_k^*(T)$.

As shown in [1], we have that $\langle F(s,T)\rangle$ is monotone in $s$ and tends to unity as $s \to \infty$; therefore, in order to compute an upper bound for $\langle F(s,T)\rangle$, we focus on $\langle F(1,T)\rangle$. With reference to Figure S1(a), in the parameter regime $r_m = 1, r_p = 0.01, r_d = 0.001, C(0) = 0.05$, there is excellent agreement between $\langle C_1(T)\rangle$ and $C(T)$, while $\max\langle F(1,T)\rangle < 1.2$ (Figure S1(b)). This indicates that the tolerance parameter $\epsilon$ should not be smaller than about 0.2, as there is no significant gain in accuracy while increasing the computational demands of the CBM.

Conversely, in the parameter regime $r_m = 1, r_p = 0.05, r_d = 0.005, C(0) = 0.05$, there is a relatively poor agreement between $\langle C_1(T)\rangle$ and $C(T)$ (Figure S1(a)), while $\max\langle F(1,T)\rangle > 1.5$ (Figure S1(b)). Therefore, a tolerance of $\epsilon > 0.5$ for uncorrelated agent pairs is too relaxed. Nevertheless, in the parameter regime $r_m = 1, r_p = 1, r_d = 0.1, C(0) = 0.05$ (Figure S1(c)), the choice of $\epsilon \in [0.1, 0.5]$ does not greatly affect $\langle s_k^*(T)\rangle$ and, consequently, $m^*$ which determines the threshold compartment size used in the CBM. Hence, we choose $\epsilon = 0.3$ in our computations of $\langle s_k^*(T)\rangle$.

We can also determine how the death rate $r_d$ affects $\langle s_k^*(T)\rangle$. With reference to Figure S1(d), employing a small to moderately-sized value of $r_d$ does not affect $\langle s_k^*(T)\rangle$ significantly. Furthermore, large values of $r_d$ are not generally biologically relevant, as proliferation assays and scratch assays focus on growing, rather than dying, populations of cells.

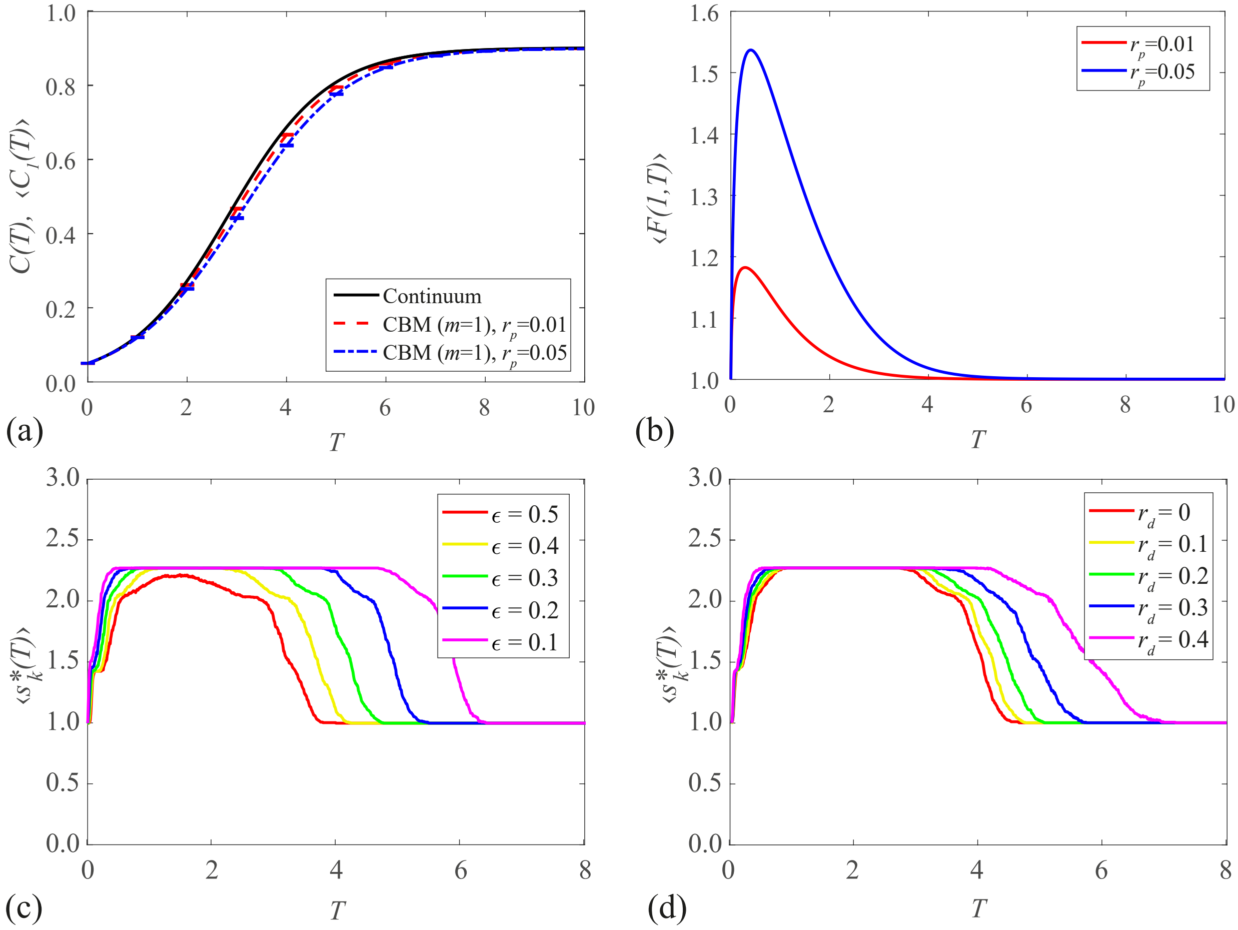


Figure S1: **Density data, correlation function, and maximum correlation distance from the CBM with $m = 1$.** The CBM uses a $120 \times 120$ square lattice with 100 realisations, $C(0) = 0.05$, $r_m = 1$, and $r_d = 0.1r_p$. (a) The CBM average agent density, $\langle C_1(T)\rangle$, and the continuum limit, $C(T)$, for $r_p = 0.01$ and $r_p = 0.05$. The maximum standard error is less than 0.0015. (b) The correlation function, $\langle F(1,T)\rangle$, for $r_p = 0.01$ and $r_p = 0.05$. (c) Comparisons of $\langle s_k^*(T)\rangle$ for various choices of $\epsilon$ with $r_p = 1$. (d) Comparisons of $\langle s_k^*(T)\rangle$ for various choices of $r_d$, with $\epsilon = 0.3$ and $r_p = 1$.

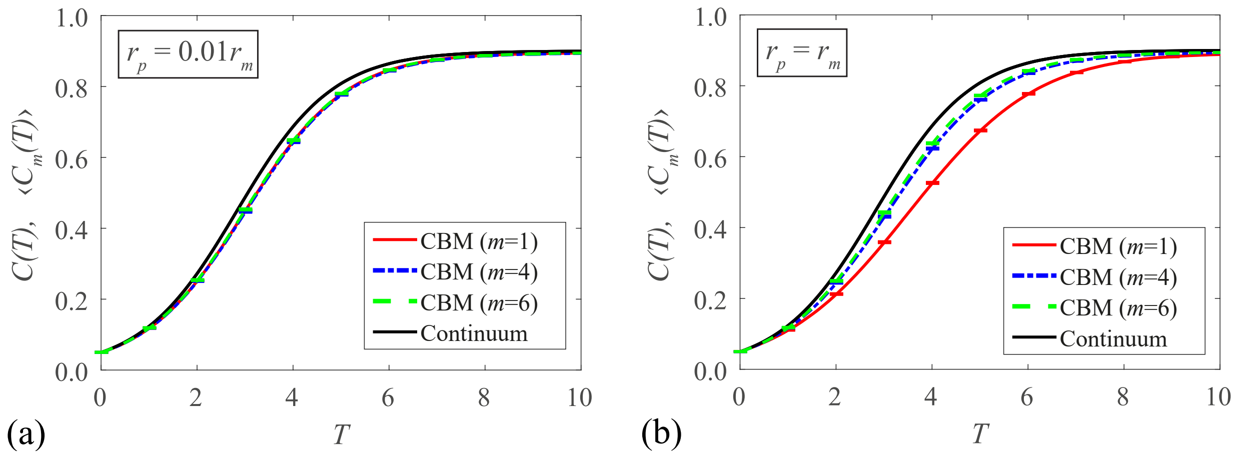


Figure S2: **Simulations of a gel proliferation assay using the CBM.** The CBM uses a $24 \times 24 \times 24$ lattice with 100 realisations, along with $C(0) = 0.05$, $r_m = 1$, and $r_d = 0.1 r_p$. The average density of agents, as determined by the CBM with $m = 1, 4$, and 6, and the solution of the continuum limit, are shown when (a) $r_p = 0.01$ (b) $r_p = 1$. The maximum standard error is less than 0.0015.

### S2 Three-dimensional simulations

Another type of assay employed in relevant cell experiments is the three-dimensional gel proliferation assay [2–4]. Gel proliferation assays are similar to two-dimensional proliferation assays, as both experiments employ cells that are initially well-mixed in growth media. However, gel proliferation assays do not produce monolayers of cells; rather, these uniformly-distributed cells proliferate throughout the three-dimensional gel in which they are seeded.

By modifying Algorithm 1 to describe a three-dimensional lattice, we can employ the CBM to describe these gel proliferation assays. To compare with the two-dimensional CBM in Section 3, which uses a $120 \times 120$ lattice, we consider a $24 \times 24 \times 24$ lattice here, as these two lattices contain roughly the same number of lattice sites. Furthermore, we use $m \times m \times m$ cubic compartments in this three-dimensional version of the CBM. With reference to Figure S2, the CBM faithfully agrees with the continuum limit in parameter regimes relevant to cell proliferation assays.

### S3 Computing the standard error of the mean

In Sections 3.1 and 3.2, error bars are included with the average CBM data describing proliferation and scratch assays. These error bars correspond to the standard error of the mean, which is calculated in one of two ways. For proliferation assays, we take the standard deviation of $C_m(T)$, denoted $\varsigma_m(T)$, using $P$ identically-prepared realisations. This allows us to calculate the standard error of the mean as a function of time:

$$\mathrm{SEM}_1(T) = \frac{\varsigma_m(T)}{\sqrt{P}}. \tag{S1}$$

For average CBM data describing scratch assays, shown in Figure 7, we record the standard error of the mean at a *fixed* time, $T_f$, as a function of $x$. This value of $T_f$ corresponds to different times displayed in the subfigures of Figure 7. To calculate this standard error, we note that for a single realisation of the CBM, we will have $Y$ observations of $C_m(x, T_f)$, which provide the vertically-averaged agent density of a single realisation of the CBM. Therefore, for $P$ identically-prepared realisations of the CBM, we have $YP$ observations of $C_m(x, T_f)$, for which we can calculate its *total* standard deviation, $\hat{\varsigma}_m(x, T_f)$. This allows us to calculate the standard error of the mean as a function of $x$ for a given $T_f$:

$$\mathrm{SEM}_2(x, T_f) = \frac{\hat{\varsigma}_m(x, T_f)}{\sqrt{PY}}. \tag{S2}$$

### S4 Varying the initial condition of the scratch assay

In Section 3.2, we examine simulations of scratch assays whose initial conditions involve some parts of the lattice being completely occupied and other parts of the lattice being completely vacant (see Figures 6 and 7 for further details concerning these initial conditions). These initial conditions represent, from a biological perspective, a region of cells at complete *confluence*, i.e. $C = 1$ in the initially occupied region of the lattice. Previous examination of scratch assays [5, 6] show that the initial cell density of a scratch assay plays a crucial role in the dynamics of the resulting moving fronts. For completeness, we also simulate scratch assays where the initial density of the initially occupied region is less than confluence, $C < 1$. To do this, each site in the non-zero region of the scratch assay simulation (for Figures 6 , 7, and S3, this region is $110 < x \leq 130$) is initially populated at random with some user-specified probability. Results in Figure S3 show that the CBM continues to capture the salient features of the scratch assay for a suitable choice of compartment size $m$. Therefore, all of the conclusions we drew in the main document for simulations of scratch assays, where the agents are initially at confluence, also apply when we consider other initial densities in scratch assays.

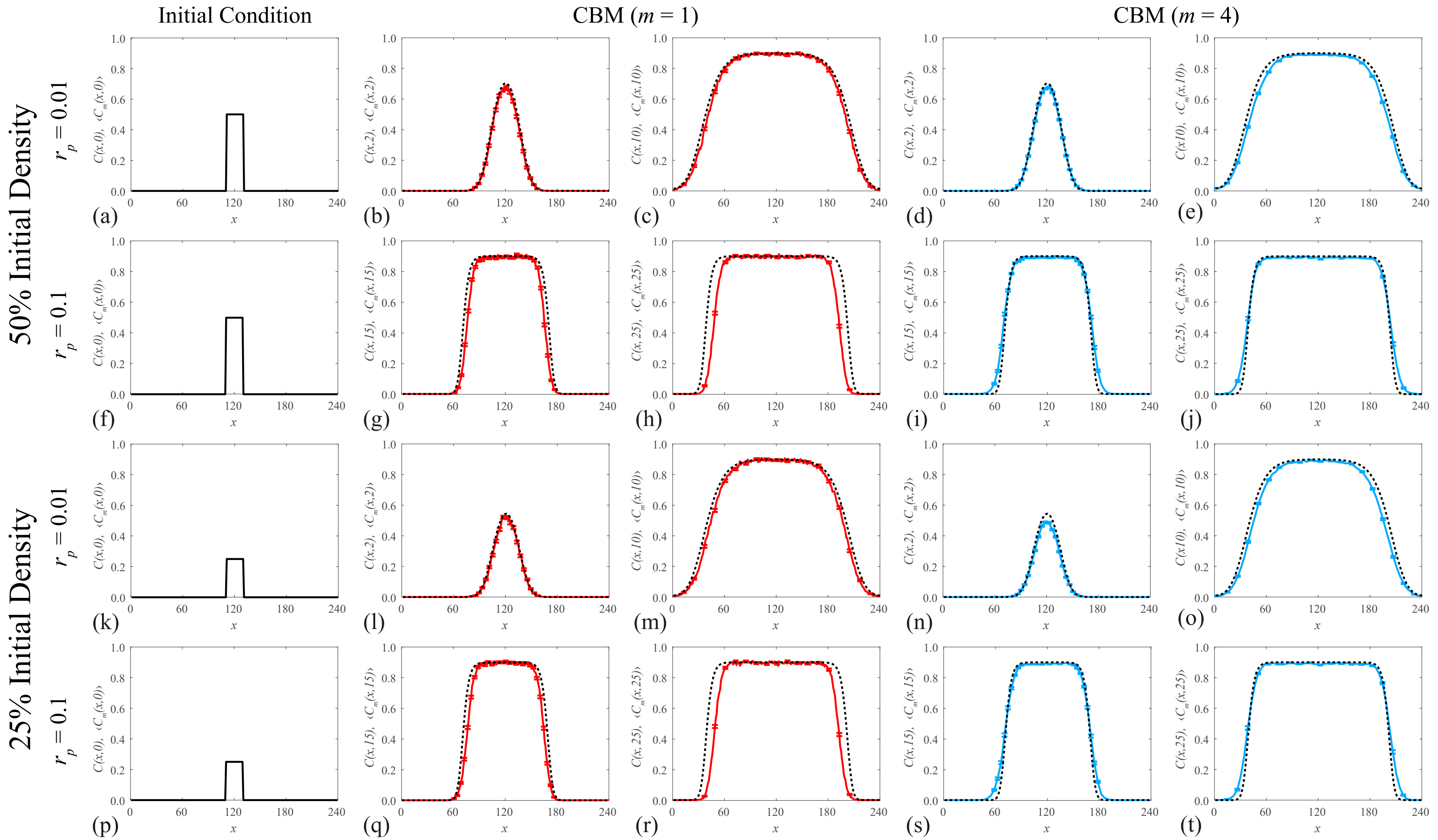


Figure S3: **Simulations of a scratch assay with the CBM with varying initial conditions.** Averaged density data from the CBM (solid colour curves) from 100 identically prepared simulations is compared with the solution of the continuum limit, Equation (11) (dashed black curves), for two different scratch assay scenarios. Results are shown for the CBM with compartment size $m = 1$ (red curves) and $m = 4$ (blue curves). In all simulations, we use a $24 \times 240$ lattice, with initial conditions shown in (a,f,k,p). The initial conditions correspond to the region $110 < x \leq 130$ being populated uniformly at random with probability (a,f) 0.5 and (k,p) 0.25. Additionally, we have $r_d = 0.1 r_p$, and $r_m = 1/m$ to keep $\hat{D}$ invariant for varying $m$. For (a)–(e) and (k)–(o), $r_p = 0.01$, with data given at (b,d,l,n) $T = 2$ (c,e,m,o) $T = 10$. For (f)–(j) and (p)–(t), $r_p = 0.1$, with data given at (g,i,q,s) $T = 15$ (h,j,r,t) $T = 25$. The maximum standard error is less than 0.011.

### S5 Biased movement

For biological experiments with chemotactic movement, cells are inclined to move in a biased direction [7–10]. One way of incorporating this phenomenon into the CBM is to bias the movement of agents by modifying Line 11 in Algorithm 1. As an example, we can bias the two-dimensional lattice examined in Section 3.2 with bias $(1+\rho)/4$ in the positive $x$-direction and bias $(1-\rho)/4$ in the negative $x$-direction, where $\rho \in [-1, 1]$. As a result, employing a positive bias $\rho$ creates a sharper front where the gradient of the agent density is positive (left front in Figure S4). Similarly, the converse is true for negative bias and we retrieve unbiased moving fronts when $\rho = 0$ (Section 3.2). As shown in [10], the continuum limit of this biased movement becomes the Fisher-Kolmogorov equation with a nonlinear advection term:

$$\frac{\partial C(x,T)}{\partial T} = C(x,T)\left(1 - \frac{C(x,T)}{K}\right) + \hat{D}\frac{\partial^2 C(x,T)}{\partial x^2} - \alpha\frac{\partial}{\partial x}[C(x,T)\,(1 - C(x,T))], \tag{S3}$$

where $\alpha = r_m\rho\Delta/(2\lambda)$. It is important to note that the advection magnitude, $\alpha$, is invariant under compartment modification. Therefore, rescaling the motility rate $r_m$ by $r_m/m$, like in Section 3.2, does not produce a compartment-invariant continuum limit when $\rho \neq 0$. Nevertheless, we can still compare the CBM with biased movement with the continuum limit (Figure S4). Like in the unbiased moving fronts case ($\rho = 0$; Section 3.2), the CBM matches this continuum limit well when $r_p/r_m \ll 1$, but only agrees well for an intermediate compartment size ($m = m^*$) when $r_p/r_m$ is no longer small.

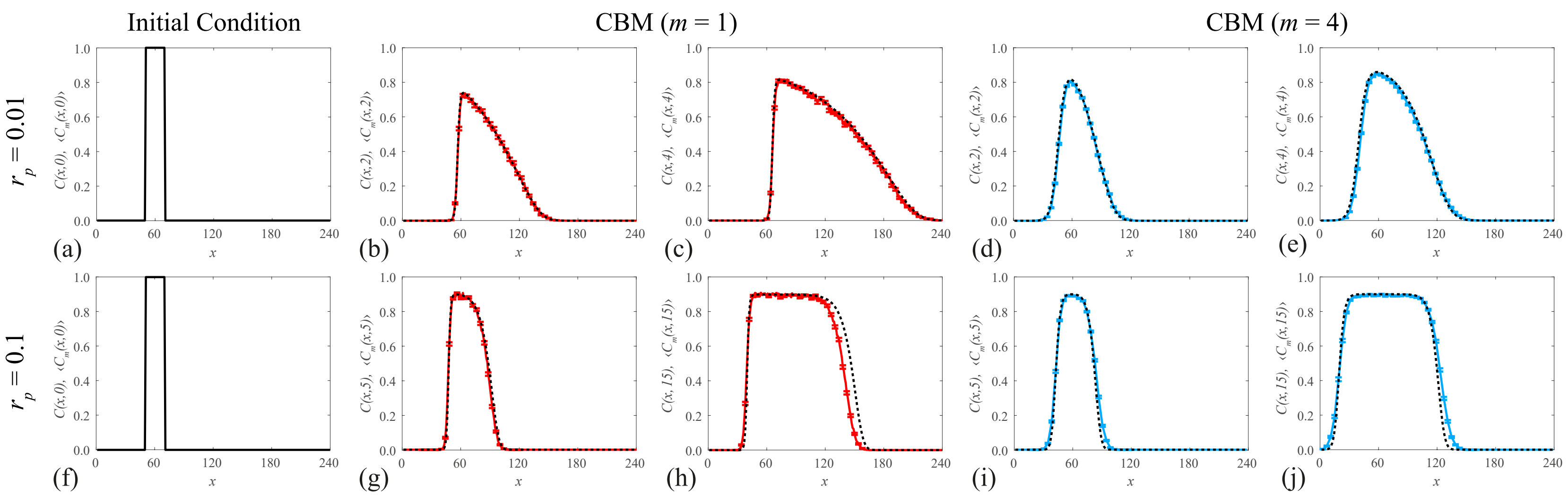


Figure S4: **Simulations of a scratch assay with biased movement using the CBM.** Averaged density data from the CBM (solid colour curves) from 100 identically prepared simulations is compared with the solution of the continuum limit, Equation (S3) (black dashed curves) with movement bias $\rho = 0.5$ in the positive $x$-direction. Results are shown for the CBM with compartment size (b,c,g,h) $m = 1$ and (d,e,i,j) $m = 4$. In all simulations, we use a $24 \times 240$ lattice, where the lattice is initially filled with agents for $50 < x \leq 70$ (a,f). Additionally, $r_d = 0.1 r_p$, and $r_m = 1/m$. Top row: $r_p = 0.01$, with data given at (b,d) $T = 2$ (c,e) $T = 4$. Bottom row: $r_p = 0.1$, with data given at (g,i) $T = 5$ (h,j) $T = 15$. The maximum standard error is less than 0.011.

### S6 Numerical solution of continuum limit

In Sections 3.2 and S5, we compare the CBM averaged data with the corresponding continuum limit (Equations (11) and (S3), respectively). To solve these PDEs numerically, we use the methods of lines in time and a second-order accurate finite difference scheme in $x$. For example, Equation (11) becomes

$$\frac{\mathrm{d}C(x_k,t)}{\mathrm{d}T} = C(x_k,T)\left(1 - \frac{C(x_k,T)}{K}\right) + \frac{\hat{D}}{\delta^2}\left[C(x_{k+1},T) - 2C(x_k,T) + C(x_{k-1},T)\right], \tag{S4}$$

where $x_{k+1} = x_k + \delta$, $x_{k-1} = x_k - \delta$, and $\delta$ is the distance between $x$ co-ordinates. The continuum limit has the domain $1 \leq x \leq I$, where $I$ is the total number of lattice sites in the $x$-direction and is defined in Section 2.1. Additionally, second-order accurate Neumann boundary conditions are used to represent the reflecting boundary conditions of the CBM:

$$C(1,T) = \frac{4}{3}C(1+\delta,T) - \frac{1}{3}C(1+2\delta,T), \tag{S5}$$

$$C(I,T) = \frac{4}{3}C(I-\delta,T) - \frac{1}{3}C(I-2\delta,T). \tag{S6}$$

Finally, we use a stiff ODE solver in time, namely, the MATLAB function `ode15s`, to achieve convergence.
